# Fecal immunoglobulin A (IgA) and its subclasses in systemic lupus erythematosus patients are nuclear antigen reactive and this feature correlates with gut permeability marker levels

**DOI:** 10.1101/2022.01.26.477918

**Authors:** Radhika Gudi, Diane Kamen, Chenthamarakshan Vasu

## Abstract

Systemic lupus erythematosus (SLE) is characterized by the production of anti-nuclear autoantibodies. Here, for the first time, we show that the abundances of gut permeability marker Zonulin and IgA1- and IgA2-subclasses are significantly higher in the fecal samples of SLE patients compared to HCs. Importantly, IgA-total, IgA1- and IgA2-subclasses from SLE patients showed higher nAg reactivity titers. Notably, we found that not only the nuclear antigen (nAg) reactive fecal IgA1:IgA2 ratio is higher in SLE patients, but also the abundance and nAg reactivity of fecal IgA and subclasses, IgA1 particularly, correlate with the fecal levels of Zonulin, which is produced primarily in the small intestine. These observations suggest that higher amounts of nAg-reactive IgA are produced, particularly, in the proximal gut as indicated by higher IgA1:IgA2 abundance and nAg reactivity ratios and correlation of these features with the Zonulin levels.

## 1. Introduction

Systemic lupus erythematosus (SLE) is an autoimmune disease that arises when abnormally functioning B lymphocytes produce autoantibodies to nuclear antigens (nAg). High levels of circulating autoantibodies and immune-complexes (IC), and IC deposition in the kidney causing glomerulonephritis are the hallmarks of SLE [1, 2]. Our preclinical studies using lupus-prone mice have shown: 1) that minor dietary deviations can alter the composition of gut microbiota and the lupus-like disease progression [3], 2) a potential contribution of pro-inflammatory immune response, including higher frequencies of plasma cells, initiated in the gut mucosa to the disease process and gender bias [4, 5], 3) higher frequencies of IgA-autoantibody producing cells in the gut mucosa, and 4) higher abundance and nAg reactivity of fecal IgA at pre-seropositive and -clinical stages [6, 7]. Importantly, higher gut permeability and microbial translocation have been implicated in the initiation and/or progression of autoimmunity in at risk subjects [8, 9].

IgA is the most abundant Ig isotype released into the gut lumen and it plays an important role in the protection against microbial infections as well as in maintaining a healthy gut microbiota [10]. In humans, two subclasses of IgA (viz: IgA1 and IgA2) are produced. In healthy human subjects, while the serum IgA antibodies are mostly IgA1, IgA antibodies secreted by intestinal B cells are composed of about 30-50% IgA1 and 50-70% IgA2 [11]. With respect to SLE, it has been shown that in addition to the differences in gut microbiota composition, the levels of total IgA in stool samples of SLE patients are relatively higher compared to that of healthy controls (HC) [12]. However, it is unknown if IgA subclasses are differently produced in the gut of lupus patients, and if their fecal IgA have higher nAg reactivity.

In the present study, we compared the abundance of pro-inflammatory marker calprotectin, permeability protein Zonulin, and IgA-total and IgA-subclasses in cross-sectional stool samples from HC subjects and recent onset SLE patients. We also examined the nAg reactivity of fecal IgA and its subclasses. We found that, in addition to relatively higher levels of calprotectin and total IgA [12], fecal levels of Zonulin, and IgA1- and IgA2-subclasses are significantly higher in SLE patients. We also found, for the first time, that fecal IgA and its subclasses, IgA1 particularly, from SLE patients have significantly higher nAg reactivity compared to that of HCs, and these features correlate with the fecal levels of gut permeability marker Zonulin. These observations suggest that fecal IgA features could serve as valuable indicators of the systemic autoimmune process in lupus patients, and potentially, at-risk subjects.

## 2. Materials and methods

### 2.1. Study subjects and samples

Cryopreserved stool sample aliquots, collected under the Institutional review board (IRB) approved protocol, were used in this study. Cross sectional stool samples from 21 adult SLE patients and 21 adult healthy controls were analyzed. Samples were either from patients with at least 4 of the 11 ACR revised criteria (for the Classification of SLE) or unaffected controls with no known personal or family history of autoimmune disease by selfreport (for the classification of HC). The original IRB protocol excluded subjects who received systemic antibiotics or commercial supplemental probiotics within the past 3 months. Those with chronic gastrointestinal disorders such as inflammatory bowel disease, irritable bowel syndrome, celiac disease, persistent, infectious gastroenteritis, colitis or gastritis, or chronic diarrhea by selfreport were also excluded. All samples were from self-identified African American women.

### 2.2. Extraction and quantitation of inflammation markers and antibodies

Stool aliquots were weighed, suspended in proportionate volume of PBS (10% suspension or 100 mg feces/ml; w/v) by breaking the fecal material using pipette tip, high speed vortexing for 1 min, and followed by shaking at 800rpm for overnight at 4°C. These suspensions were centrifuged at 14,000 rpm for 10 min, upper 2/3 portion of the supernatants were separated and used in ELISA assays for estimating Calprotectin, Zonulin, IgA, IgA1 and IgA2 levels. Calprotectin levels were determined in optimally diluted (1:20) extracts using Human-S100A8/S100A9 Heterodimer DuoSet-ELISA kit from R&D Systems. Zonulin levels were determined in similarly diluted extracts using human Zonulin Stool ELISA kit from ALPCO. For detection of IgA, IgA1 and IgA2, in-house sandwich ELISAs with purified, commercially available antibodies and standards for generating standard curves were employed.

For IgA, IgA1 and IgA2 sandwich ELISAs, purified anti-mouse IgA, IgA1 and IgA2 monoclonal antibody (0.1μg/well in 50μl) coated wells were incubated with optimum dilution of samples for 2h. Wells were incubated further with biotin-linked polyclonal anti-mouse IgA, IgA1 and IgA2 antibody for 1h, and finally with streptavidin-HRP for 30 min, before developing the reaction using TMB/H_2_02 substrate and read at 450nm. Optimum dilutions of extracts were 1:3000 for IgA, 1:500 for IgA1, and 1:1000 for IgA2. Purified mouse IgA, IgA1 and IgA2 (Athens Research and Southern Biotech) were used in each plate for generating respective standard curves to calculate the concentration in 100mg stool samples. These values were then converted to respective antibody concentration per gram of stool with a multiplication factor of 10.

### 2.3. Determination of nAg reactivity titer of fecal antibodies

Antibody titers against nAgs (nucleohistone and dsDNA) in aforementioned fecal extracts were determined by employing inhouse indirect ELISA [3, 5] with necessary modifications. Briefly, 0.5μg/well of nucleohistone (Sigma-Aldrich) or dsDNA from calf thymus (Sigma-Aldrich) was coated as antigen (carbonate buffer or DNA-coating reagent from Invitrogen respectively), overnight, onto ELISA plate wells. These wells were then incubated with serial dilutions of the fecal extracts (starting at 1:20 dilution) for 2 hours and followed by 1 hour incubation with biotin-linked anti-human IgA, IgA1, and IgA2 antibody. Streptavidin-HRP was added to the well for 30 min, before developing the reaction using TMB/H_2_0_2_ substrate and the plates were read at a wavelength of 450 nm. Highest dilution of the sample that produced an OD value of ≥0.05 above background value was considered as the nAg reactive titer of 100 mg feces. These titer values were then converted to per gram nAg reactivity titer with a multiplication factor of 10.

### 2.4. IFA staining of Hep2 cells to detect ANA

Kallestad Hep-2 Complete Kit (Bio-rad) was used to detect ANA reactive fecal IgA following the kit instructions. 1:40 dilution of fecal extracts was employed. Anti-human IgG-FITC from the kit was replaced with anti-human IgA-Alexa flour 488 reagent as the secondary antibody to detect fecal IgA antibody. Relative fluorescence intensities were graded arbitrarily on a 0-4 scale using a fluorescent microscope.

### 2.5. Statistical analysis

Mann-Whitney test using GraphPad Prism was employed to calculate *p*-values. A *P* value of ≤0.05 was considered statistically significant.

## 3. Results

### 3.1. Levels of fecal proinflammatory/permeability marker and IgA and its subclasses (IgA1 and IgA2) are higher in SLE patients

Extracts were prepared from frozen stool aliquots of 21 new onset SLE patients and 21 HCs and subjected to quantitative ELISA to determine the levels of calprotectin, zonulin, IgA, IgA1 and IgA2. As observed in **Fig. 1A**, protein levels of the pro-inflammatory marker calprotectin were relatively higher, albeit not significant statistically, in stool samples from SLE patients compared to HC. **Fig. 1B** shows that fecal levels of the gut permeability marker zonulin were significantly higher in SLE patient samples compared to HC samples. As reported before [12], total IgA levels were also significantly higher in fecal samples of SLE patients (**Fig. 1C**). Importantly, the levels of both IgA subclasses, IgA1 and IgA2, were significantly higher in SLE patient samples compared to that of HCs (**Fig. 1D & 1E**). To assess if SLE patients, compared to HCs, show different degrees of IgA1 and IgA2 secretion in the gut, IgA1:IgA2 ratios were calculated for each sample. As observed in **Fig. 1F**, although not significant statistically, IgA1:IgA2 ratios were relatively higher in SLE patients compared to controls. These results show that while the levels of overall IgA and both subclasses are higher in the stool samples of SLE patients, these patients appear to release relatively higher amounts of IgA1 than IgA2 into the gut lumen as compared to HCs.

**Figure 1:**
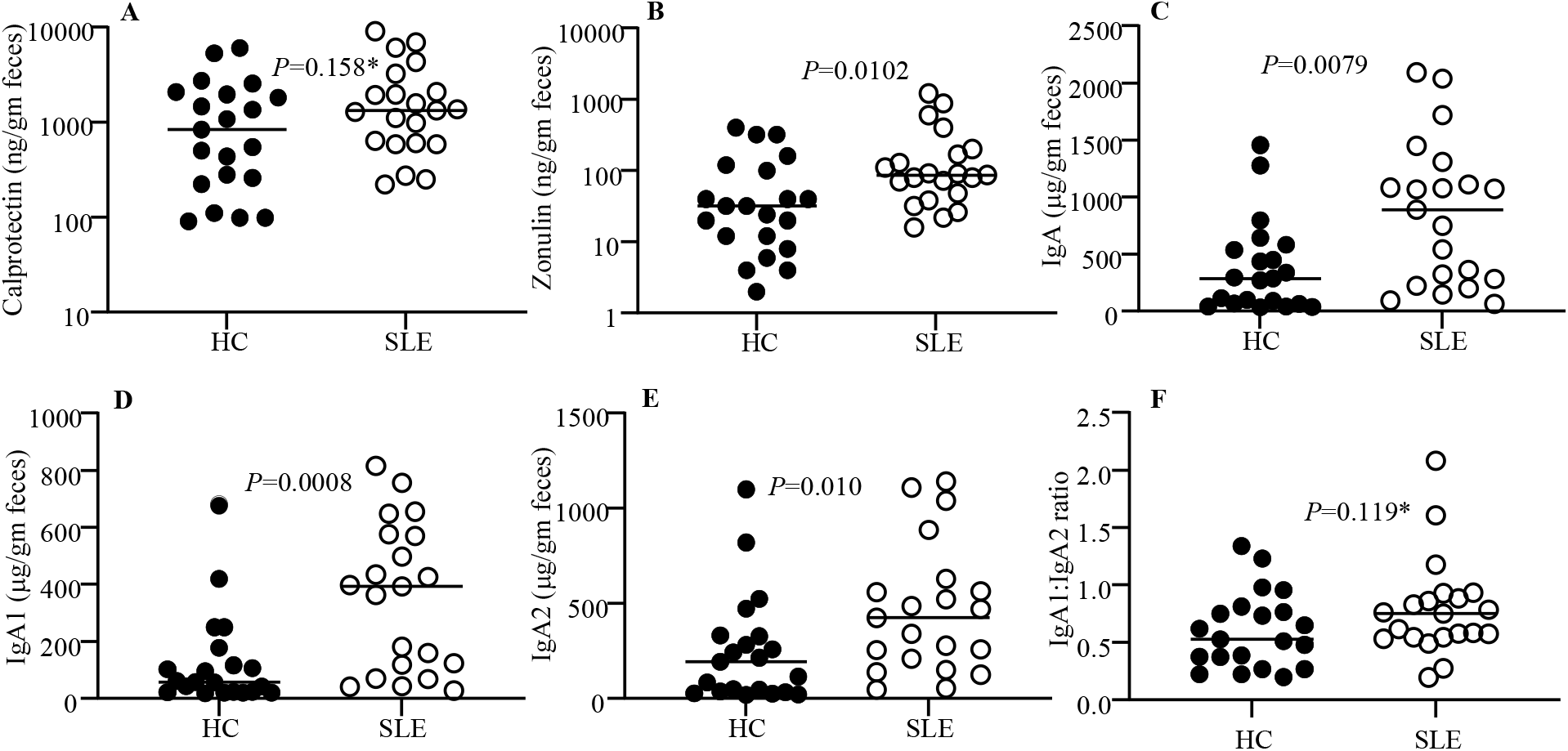
Levels of fecal proinflammatory/permeability marker, and IgA and its subclasses (IgA1 and IgA2) in HC subjects and SLE patients. Optimally diluted PBS extracts of stool samples from HC subjects (n=21) and SLE patients (n=21) were tested for protein levels of calprotectin **(A)**, Zonulin **(B)**, IgA **(C)**, IgA1 **(D)** and IgA2 **(E)** as described under materials and methods section. IgA1: 1gA2 ratios are also shown **(F)**. *P*=values by Mann-Whitney test.

### 3.2. Fecal IgA and its subclasses show greater nAg reactivity

To determine if fecal IgA of SLE patients have higher nAg reactivity, dsDNA and NH reactive IgA titers of extracts prepared from stool samples were determined by ELISA. As shown in **Fig. 2A**, both dsDNA and NH reactive IgA titers were significantly higher in the samples of SLE patients compared to that of HCs. Further, Hep2 cell substrate-based immunofluorescence assay showed similarly higher nAg reactivity by fecal IgA of SLE patients **(Fig. 2B)**. Examination of the nAg reactivities of IgA subclasses revealed that the dsDNA and NH reactive IgA1 and IgA2 antibody titers are higher in SLE patients compared to HCs **(Fig. 2C)**. Importantly, differences in the dsDNA and NH reactive titers of SLE patients and HCs were more pronounced with IgA1 than IgA2. Therefore, dsDNA and NH reactive fecal IgA1: IgA2 ratios were determined. As observed in **Fig. 2D**, nAg reactive IgA1:IgA2 ratios were significantly higher in samples from SLE patients than that from HCs. Overall, these results show that the nAg reactivity of fecal IgA is higher with SLE patient samples, and this is more pronounced with IgA1.

**Figure 2:**
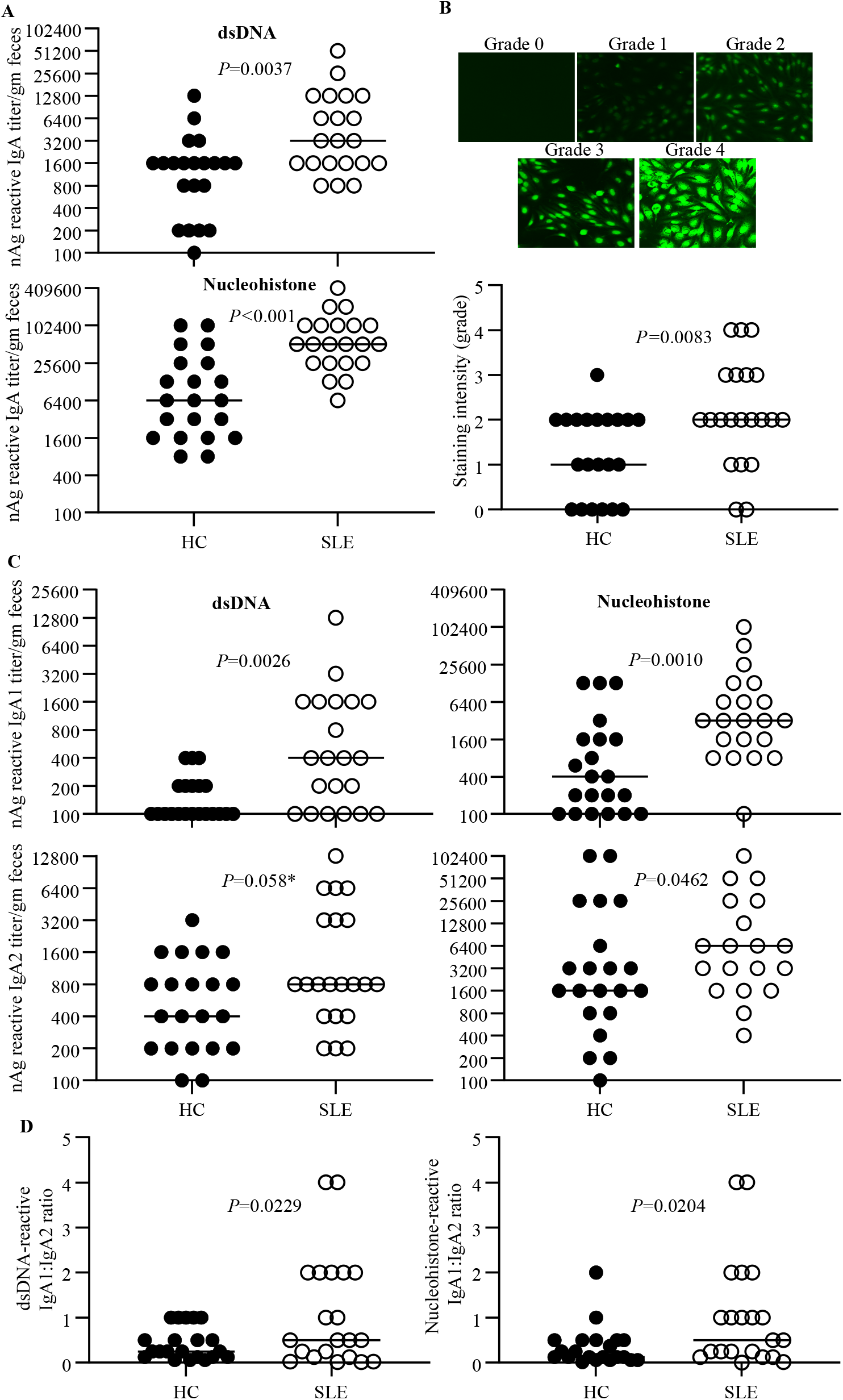
nAg reactivity of fecal IgA and its subclasses. PBS extracts of stool samples from HC subjects (n=21) and SLE patients (n=21) were examined for nAg reactivity as described under the Materials and methods section. **A)** nAg reactivity of total IgA was determined with dsDNA and NH(nucleohistone) coated plates, 2-fold serial dilutions of the extracts, and anti-human-IgA-biotin as the secondary antibody. **B)** Hep-2 slides were stained using 1:40 dilution of fecal extracts and anti-human IgA-Alexa flour 488 reagent and imaged under a fluorescent microscope. Relative fluorescence intensities were graded arbitrarily on a 0-4 scale. Representative images (upper panel) and overall intensities (lower panel) are shown. **C)** nAg reactivity of total IgA was determined as described for panel A using anti-human-IgA1- or IgA2-biotin as the secondary antibody. **D)** dsDNA reactive IgA1: 1gA2 ratios are shown. *P*=values by Mann-Whitney test.

### 3.3. Zonulin levels correlate with fecal IgA abundance and nAg reactivity

Since the levels of Zonulin in the fecal samples were significantly higher in SLE patients compared to HCs, we examined if the abundance and nAg reactivity of fecal IgA correlate with the protein levels of this gut permeability protein in each group. Interestingly, while Zonulin levels in the fecal samples of both HC and SLE groups showed significant correlation with total IgA and IgA1 abundances (**Fig. 3A**) nevertheless, samples from SLE group showed an overall better correlation of Zonulin levels with both subclasses of IgA as well as total IgA levels. Importantly, while the Zonulin levels of samples from both HC and SLE groups showed significant correlation with dsDNA reactivity of total IgA, Zonulin levels of samples from SLE group, but not HC group, correlated significantly with nAg reactivity of IgA1. Overall, these observations suggest that Zonulin dependent higher gut permeability, particularly in the proximal gut, of SLE patients, and the associated pro-inflammatory events could be contributing to the higher abundance and nAg reactivity of fecal IgA.

**Figure 3:**
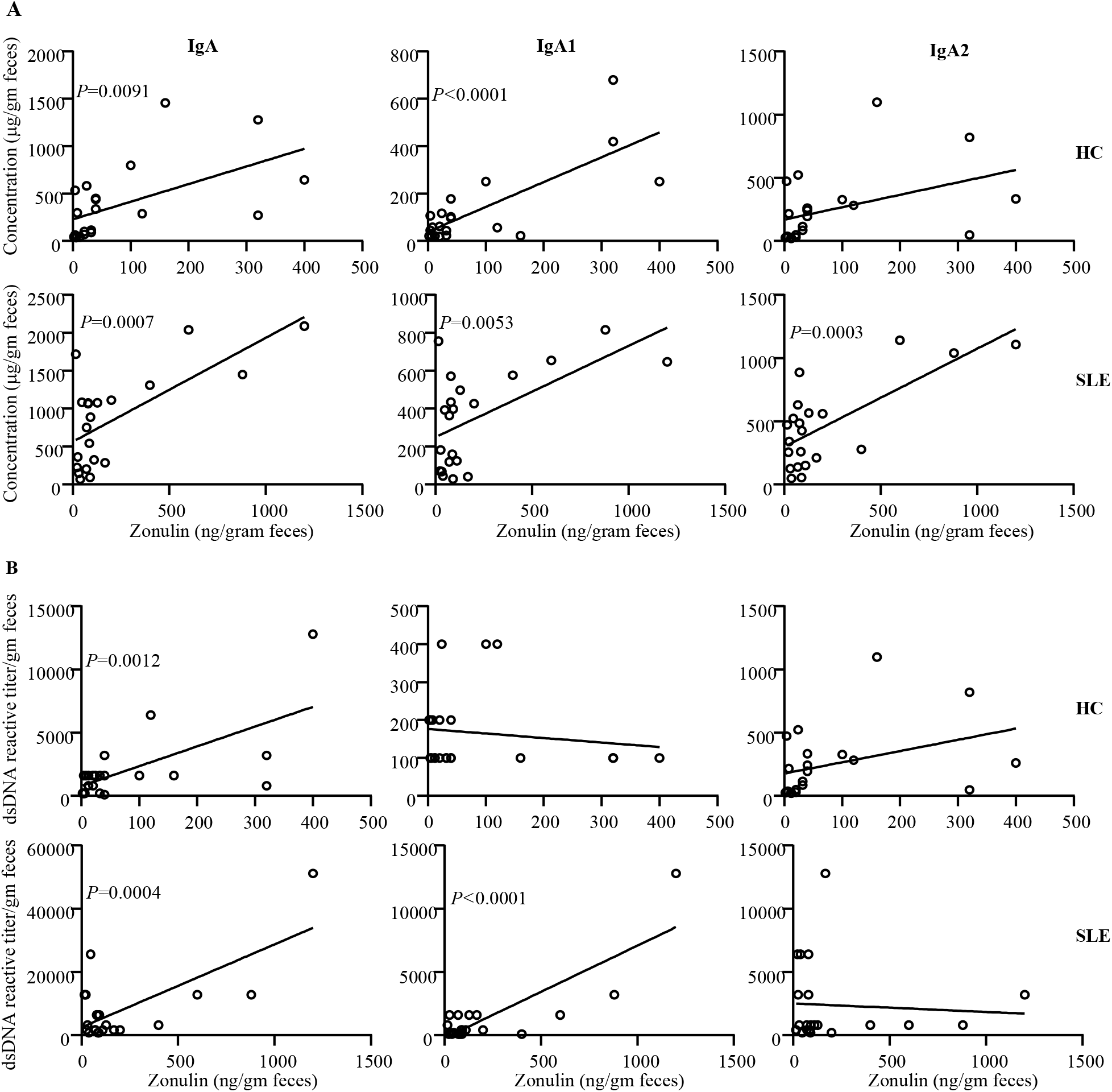
Zonulin levels correlate with the fecal IgA abundance and nAg reactivity. **A)** Fecal IgA, IgA1 and IgA2 concentrations of samples from HC subjects and SLE patients were correlated with respective fecal Zonulin concentration values by Pearson’s correlation approach. **B)** dsDNA reactivity titers of fecal IgA, IgA1 and IgA2 of HC subjects and SLE patients were correlated with respective fecal Zonulin concentration values by Pearson’s correlation approach.

## 4. Discussion

Our recent studies have shown that in a mouse model of spontaneous autoimmune lupus, pro-inflammatory phenotype of the gut mucosa at as early as juvenile age correlates with the gender bias observed in lupus-like disease[4, 5]. High levels of a pro-inflammatory factor, calprotectin were detected in the fecal samples of SLE patients[12]. On the other hand, a role of the gut permeability protein Zonulin, which is produced primarily in the small intestine, in autoimmune and inflammatory conditions has also been described[13]. However, whether Zonulin is released at higher levels in the gut lumen of SLE patients is not known. Our observations that Zonulin levels are significantly higher in the fecal samples of SLE patients, suggest that SLE patients, and potentially at-risk subjects, may have higher gut permeability due to the higher levels of Zonulin protein being produced in the gut.

Pro-inflammatory events of the gut mucosa can contribute to the production and release of antibodies, IgA particularly, in the gut lumen. While a recent report has shown that fecal IgA levels are higher in SLE patients compared to HCs [12], we reported that fecal IgA of lupus-prone mice have higher nAg reactivity long before the clinical disease onset [6]. In agreement with the previous report [12], we found that stool samples from majority of the 21 SLE patients tested in our study had higher amounts of soluble IgA than HCs. Importantly, however, we report that not only both IgA1 and IgA2 subtypes levels are higher in the fecal samples of SLE patients, but also IgA antibodies in general, and IgA1 in particular, have significantly higher nAg reactivity compared to that of HCs.

IgA produced in the human intestine is comprised of two subclasses, IgA1 and IgA2 [10]. While human serum contains predominantly IgA1, feces contains both IgA1 and IgA2 with higher amounts of the latter. This is because while plasma cells in the small intestine secrete mostly IgA1, the proportion of IgA2 production increases from the duodenum and this subclass is produced at profoundly higher levels in the colon. Although higher B cell function and antibody production occurs in SLE patients, it is not known whether IgA1 and IgA2 subtypes are produced disproportionately in the gut mucosa of SLE patients compared to HCs. Our observations that while the overall abundances of both IgA1 and IgA2 are higher in the fecal samples of most SLE patients, IgA1:IgA2 ratio is higher in SLE patients compared to HCs suggesting a relatively higher activation of IgA producing B cells in the upper digestive tract of SLE patients, compared to the distal gut.

Autoantibody response against nucleic acids and nuclear proteins including histones is the key feature of lupus in humans and rodent models. Our observation that fecal IgA, both IgA1 and IgA2 subclasses, of SLE patients have higher nAg reactivity suggests that IgA response to gut microbes could be contributing to the autoimmune process in SLE. Of note, IgA produced in the gut mucosa are mostly poly-reactive and previous studies have recognized some degree of self-antigen reactivity of fecal IgA and/or B cell clones from the gut mucosa of healthy individuals [14]. Hence it was not surprising that fecal IgA of many HCs also showed nAg reactivity. Nevertheless, SLE patients have significantly higher nAg reactive fecal IgA than the HCs. Higher IgA production and nAg reactivity of fecal IgA, particularly IgA1, in SLE patients could also be possibly associated with higher gut permeability associated inflammation in SLE patients. Since IgA1 and Zonulin are produced mainly in the small intestine [13, 15], higher correlation of Zonulin levels with IgA and IgA1 abundance and nAg reactivity in SLE patients does support this notion.

In conclusion, we found, for the first time, that both IgA1 and IgA2 levels are significantly higher in SLE patients compared to HCs. Most importantly, we also found that fecal IgA, both IgA1 and IgA2 subclasses, of SLE patients are more nAg reactive compared to that of HC subjects. Furthermore, fecal IgA1:IgA2 abundance and nAg reactivity ratios are higher in SLE patient samples compared to that of HCs. With respect to potential mechanisms, higher Zonulin-dependent gut permeability and the associated inflammation of proximal intestinal regions could be contributing to these higher fecal IgA and nAg reactivity in SLE patients. Our observations suggest that, in addition to the higher abundance and nAg reactivity of fecal IgA and its subclasses, particularly IgA1, in association with Zonulin levels could serve as a biomarker and be indicative of the severity of systemic autoimmunity in SLE patients. In this regard, it would be informative to determine if fecal IgA and subclass features and Zonulin alter during active disease flares in SLE patients. Furthermore, although additional studies using longitudinal samples from subjects who are susceptible to SLE are needed, our current observations in the context of our preclinical studies [6, 7] suggest the potential biomarker value of fecal IgA features alone or along with Zonulin levels, in predicting the ongoing autoimmune process, if any, in at-risk subjects.

## Acknowledgments

This work was supported by internal funds from MUSC, National Institutes of Health (NIH) grants R21AI136339 and R01AI138511 to CV and R21AR067459 and K24AR068406 to DK. R.G. performed experiments and reviewed the paper, DK acquired clinical samples under IRB approved protocols, and C.V. designed the study, performed the experiments, and wrote the paper. Dr. Vasu is the guarantor of this work and, as such, had full access to all the data and takes responsibility for the integrity and accuracy of the data and analysis.

## References

[1] R. Gualtierotti, M. Biggioggero, A. E. Penatti, P. L. Meroni. Updating on the pathogenesis of systemic lupus erythematosus. Autoimmun Rev, 2010 10:3–7.

[2] L. Li, C. Mohan. Genetic basis of murine lupus nephritis. Semin Nephrol, 2007;27:12–21.

[3] B. M. Johnson, M. C. Gaudreau, M. M. Al-Gadban, R. Gudi, C. Vasu. Impact of dietary deviation on disease progression and gut microbiome composition in lupus-prone SNF1 mice. Clin Exp Immunol, 2015;181:323–37.

[4] B. M. Johnson, M. C. Gaudreau, R. Gudi, R. Brown, G. Gilkeson, C. Vasu. Gut microbiota differently contributes to intestinal immune phenotype and systemic autoimmune progression in female and male lupus-prone mice. J Autoimmun, 2020;108:102420.

[5] M. C. Gaudreau, B. M. Johnson, R. Gudi, M. M. Al-Gadban, C. Vasu. Gender bias in lupus: does immune response initiated in the gut mucosa have a role? Clinical and experimental immunology, 2015;180:393–407.

[6] W. Sun, R. R. Gudi, B. M. Johnson, C. Vasu. Abundance and nuclear antigen reactivity of intestinal and fecal Immunoglobulin A in lupus-prone mice at younger ages correlate with the onset of eventual systemic autoimmunity. Sci Rep, 2020;10:14258.

[7] R. Gudi, S. Roy, W. Sun, C. Vasu. Preclinical stage abundance and nuclear antigen reactivity of fecal Immunoglobulin A (IgA) varies among males and females of lupus-prone mouse models. Immunology, 2022.

[8] A. Lerner, T. Matthias. Changes in intestinal tight junction permeability associated with industrial food additives explain the rising incidence of autoimmune disease. Autoimmun Rev, 2015;14:479–89.

[9] S. Manfredo Vieira, M. Hiltensperger, V. Kumar, D. Zegarra-Ruiz, C. Dehner, N. Khan et al. Translocation of a gut pathobiont drives autoimmunity in mice and humans. Science, 2018;359:1156–61.

[10] G. P. Donaldson, M. S. Ladinsky, K. B. Yu, J. G. Sanders, B. B. Yoo, W. C. Chou et al. Gut microbiota utilize immunoglobulin A for mucosal colonization. Science, 2018;360:795–800.

[11] J. M. Woof, M. W. Russell. Structure and function relationships in IgA. Mucosal Immunol, 2011;4:590–7.

[12] D. Azzouz, A. Omarbekova, A. Heguy, D. Schwudke, N. Gisch, B. H. Rovin et al. Lupus nephritis is linked to disease-activity associated expansions and immunity to a gut commensal. Ann Rheum Dis, 2019;78:947–56.

[13] A. Fasano. Zonulin, regulation of tight junctions, and autoimmune diseases. Ann N Y Acad Sci, 2012;1258:25–33.

[14] J. J. Bunker, A. Bendelac. IgA Responses to Microbiota. Immunity, 2018;49:211–24.

[15] A. Tripathi, K. M. Lammers, S. Goldblum, T. Shea-Donohue, S. Netzel-Arnett, M. S. Buzza et al. Identification of human zonulin, a physiological modulator of tight junctions, as prehaptoglobin-2. Proceedings of the National Academy of Sciences of the United States of America, 2009;106:16799–804.

